# Plural breeding among unrelated females and other insights on complex social structure in the cooperatively breeding Variegated Fairywren

**DOI:** 10.1101/2023.03.01.530581

**Authors:** Jordan Boersma, Derrick J. Thrasher, Joseph F. Welklin, Daniel T. Baldassarre, Michael S. Webster

**Affiliations:** Department of Neurobiology and Behavior, Cornell University, USA; Cornell Lab of Ornithology, Ithaca, NY, USA; Department of Biology, University of Nevada, Reno, Nevada, USA; Department of Biological Sciences, SUNY Oswego, Oswego, NY, USA

## Abstract

Cooperatively breeding species vary widely in degree of social complexity, and disentangling relationships among group members can reveal the costs and benefits of cooperation. Here, we explore the social system of a relatively unstudied cooperatively breeding bird, the Variegated Fairywren (*Malurus lamberti*), and explore how social complexity and group dynamics may affect cooperation and conflict. We used a combination of field-based population monitoring and detailed social association observations to determine group membership annually across four breeding seasons (2014 – 2017), and used a ddRAD-seq genotyping method to determine genetic relationships within social groups. Social groups ranged in size from 2 – 8 individuals and nearly half of all social groups had multiple adult individuals of both sexes. Approximately two-thirds of those groups exhibited plural breeding, in which multiple females within the same social group nested individually on the same territory. Genetic relationships were diverse across social groups, and many consisted of a combination of relatives and non-relatives of each sex. Notably, although related females often were present within a social group, co-breeding females in the same social group were never closely related to each other. Given extensive variation in relatedness among group members, cooperation in the Variegated Fairywren is likely maintained by a combination of direct and indirect fitness benefits.

## Introduction

Cooperative breeding – wherein multiple individuals aid in the rearing of offspring – is widespread across taxa (Koenig and Dickinson 2016). Many cooperatively breeding animals live in family groups consisting of a single breeding pair and related non-breeding helpers (Rubenstein and Shen 2009; Koenig and Dickinson 2016). In these cases, group members receive kin selected benefits from cooperating with close relatives (Dickinson 2004; Pizzari and Gardner 2012). However, factors such as turnover of breeders and extra-pair paternity (EPP) can complicate patterns of genetic relatedness among individuals within a group, potentially leading to reduced inclusive fitness benefits (Hamilton 1963; Hamilton 1964; Bourke 2014). Nearly half of cooperatively breeding bird species live in social groups containing some combination of unrelated and related individuals (Riehl 2013). Yet the vast majority of studies of cooperative breeding have focused on species living in family groups (Hatchwell and Komdeur 2000; Shen *et al*. 2017a), and studies of species that live in more genetically complex groups are far less common (Painter *et al*. 2000; Clutton-Brock 2009; Riehl 2013), thus constraining our understanding of the evolution of cooperation.

When genetic relationships among cooperative group members are mixed, kin-selected benefits are diminished for some group members, and other benefits or constraints are likely necessary to explain cooperation among non-relatives (Painter *et al*. 2000; Dickinson 2004; Clutton-Brock 2009; Kingma *et al*. 2011; Carter and Wilkinson 2015). For instance, group members may benefit from the opportunity to fill vacated breeding roles (Cockburn *et al*. 2008; Shen *et al*. 2017a), which is more likely to occur in groups containing multiple breeding pairs. Cooperation in this species could also allow groups to buffer against detrimental effects of inhabiting unpredictable environments (Rubenstein and Lovette, 2007) and ward off the abundant and diverse suite of brood parasites in Australia (Feeney *et al*. 2013). Cooperatively-breeding species that have both unrelated and related individuals cooperating are particularly interesting because multiple types of benefits of cooperation may operate simultaneously (Dickinson 2004; Clutton-Brock 2009; Rubenstein *et al*. 2016).

The Australasian Fairywrens (*Malurus* spp.) have served as model systems for studies of cooperative breeding and complex social behavior for several decades (Buchanan and Cockburn 2013; Cockburn *et al*. 2013; Joseph *et al*. 2013). Many Fairywren species live in simple family groups, with a breeding pair and several helping (often male) auxiliaries (Rowley & Russell, 1997). However, genetic relatedness in these species can be complicated by high EPP rates (Webster *et al*. 2004; Varian-Ramos and Webster 2012; Cockburn *et al*. 2013; Brouwer *et al*. 2017) and rapid replacement of breeders when a vacancy opens (Varian-Ramos and Webster 2012). In some Malurids, female auxiliaries also serve as non-breeding helpers (Russell and Rowley 2000) and are often daughters that delay dispersal and help in their natal group. Independent reproduction by multiple pairs within social groups (i.e. plural breeding) is uncommon in Fairywrens (but see Rowley et al. 1989; Buchanan and Cockburn 2013), but appears to be more likely when immigrant females join established groups (Brouwer *et al*. 2011; Johnson and Pruett-Jones 2018). Thus, high EPP rates and immigration can lead to complex patterns of relatedness within social groups, likely affecting the relative costs and benefits of group membership.

Here we examine group composition and social dynamics in the Variegated Fairywren (*Malurus lamberti*) of eastern Australia, a member of the “chestnut-shouldered clade” of Fairywrens, recently split from its sister species, the Purple-backed Fairywren (McLean et al. 2017b; McLean et al. 2017a). Natural history studies for this species have shown that social group size can vary substantially, and female auxiliaries of unknown reproductive status have been reported (Rowley and Russell 1997). Despite being a rather conspicuous and common species, little is known about its social system, particularly genetic relationships of group members and breeding behavior. We intensively monitored a population of variegated-Fairywrens near Brisbane, Queensland to study the social dynamics and genetic relationships of this complex cooperatively breeding species. We used detailed field observations to describe overall associative behavior and territoriality, determined group size and composition, and designated social statuses within groups. Additionally, we used a unique panel of single nucleotide polymorphisms (SNPs) to determine relatedness of adults within social groups. Together, these approaches allowed us to show that this species exists in especially complex social groups, often containing multiple breeding pairs and helpers, with relationships among group members varying from completely unrelated to parent-offspring relationships.

## Methods

### Study Population and General Field Methods

We studied a population of Variegated Fairywrens (*M. lamberti*) on the shore of Lake Samsonvale (27°160 S, 152°410 E), 30 km northwest of Brisbane, Queensland, Australia. We collected data on color-banded birds throughout the breeding season each year, typically August – January, from 2014 – 2017. The predominant habitats at our study site include subtropical grassland, eucalypt plantations, and dry eucalypt forest with secondary growth. Variegated Fairywrens at our study site largely occupied areas of secondary growth but were also present to a lesser extent in other habitat types, such as open grassland and lake margins. We captured birds using targeted mist-netting, occasionally using playback of distress calls as a lure. We banded each adult with a unique combination of three plastic color bands and an Australian government issued (ABBBS) aluminum band for individual identification. We collected a jugular blood sample of 50 – 80μl from all individuals for subsequent genetic analyses (Baldassarre & Webster, 2013), as well as standard morphometric measurements including mass, tarsus and tail length, wing chord, and bill measurements. When possible, we determined the age and sex of each captured individual using plumage characteristics (i.e. nuptial plumage, Johnson and Pruett-Jones 2018), ossification of the skull (Lindsay *et al*. 2009), and physical indicators of reproductive status (i.e. brood patch or cloacal protuberance). Throughout each breeding season we systematically monitored the population by assessing group membership, affiliative behaviors, and breeding activity. The geography of the field site (i.e., bordered on multiple sides by water) also afforded close monitoring of dispersal and movement of individuals born into the population, and immigrants entering the population.

### Social Group Composition

Social group membership and territories were determined during routine monitoring, using repeated observations of individuals across the study site. Within the first few weeks of each field season, we defined social groups as any aggregation of two or more individuals that were present in the same area on more than three occasions, and that were observed engaging in associative behavior (e.g., allopreening, foraging in close proximity) with each other or in coordinated defensive behaviors (i.e., territoriality) against conspecifics. Repeated observations in each season allowed us to confirm our designations, assess any changes in social group composition, and determine female breeding status. Although most groups were stable within a breeding season, some did undergo changes in size and/or composition. For these cases, to avoid pseudoreplication, we considered the group’s most complex arrangement of adult birds within each season for subsequent analyses, defined as being the largest in size, having multiple females (breeding or non-breeding), and/or having multiple breeding females. In most cases, an individual’s most complex group was its first social group of the season.

During the 2016 field season we conducted additional systematic observations of associations among individuals and used social network analysis to corroborate our routine monitoring group assignments described above. To identify groups using social networks, we conducted 25 minute focal follows of groups, collecting data on which birds were associating every five minutes, resulting in 6 sampling points per observation. Birds were considered associating if they were within a 30-meter area and moving and vocalizing in a coordinated manner. We followed most social groups for at least three 25-minute observations (mean = 5.82 observations/group) during the breeding season. We constructed a social network using the gambit of the group method, considering any individual associating in a sampling point to be associating in the network. We built the network using the simple ratio index (SRI) in the R package ‘asnipe’ (Farine 2013) and then removed individuals that were seen fewer than 7 times, meaning an individual had to be seen in at least two observations to be included in the network to make sure we had accurately assessed each individual’s social relationships.

We identified groups in the social network using a dendrogram method (see Welklin et al. 2023). To summarize, we created a dendrogram using the UPGMA method, then searched for the bifurcation point in the dendrogram that was associated with the highest average silhouette width when the dendrogram was cut at that point. Silhouette width is a clustering quality score that compares the distances between nodes (individuals) within a cluster (social group) to the distance to the next-closest cluster. A score close to 1 indicates a network with distinct clusters and a score close to 0 indicates a very uniform network. Individuals alone in solo “groups” were not included in subsequent analyses as we never observed floaters in our population, and it was more likely that other group members associated with these individuals were not observed often enough to be included in the network. We compared the structure of these network-defined groups to that of data-stream permuted networks to test whether the observed groups were more structured than expected by random chance (Welklin *et al*. 2023). We compared the membership of field-defined and network-defined social groups by calculating the percentage of within-group dyads that were in the same groups across the two different methods.

### Monitoring Reproduction

We found most or all nests and intensively monitored nesting attempts for all social and breeding groups throughout each breeding season. Females within each group were designated as “breeding” if they were observed actively engaging in nest-building, incubation or brooding, or if they were captured with an active (i.e. defeathered and vascularized) brood patch. In Fairywrens, only the breeding female builds the nest and incubates the eggs (Schodde 1982; Rowley and Russell 1997). Males that attended to the female closely throughout nest building and early nestling rearing were designated as breeding social mate(s) of the breeding female. Non-breeding individuals within a group were those without their own nest (i.e., females that never built a nest and males that were not associated with a nest-building female). We determined nest fate by checking each nest once every three days until failure or fledging. When nestlings reached six days of age, they were banded with an aluminum ABBBS band and we took a small tarsal blood sample for genetic analysis of parentage. Nestlings that survived to day nine were banded with a unique combination of color bands for later identification.

### SNP Genotyping and Pairwise Relatedness

We used a diverse set of single nucleotide polymorphisms (SNPs) to address questions about genetic relatedness among individuals in social groups. The SNP panel was derived using a double-digest restriction-site associated DNA sequencing approach (ddRAD-seq) described in Thrasher et al. (2018). A total of 858 individuals were sequenced across five sequencing runs using an Illumina HiSeq 2500 with single-end reads. Following sequencing, we used a *de novo* assembly for subsequent SNP calling. After filtering for missing data, depth of coverage, and minimum allele frequency, the SNP panel consisted of 358 unique markers (see Thrasher et al. 2018 for filtering metrics).

We used the package, “RELATED” (Pew et al., 2015), in R version 3.5.3 (R Core Team 2019) to estimate pairwise relatedness (*r*) for all adults within each social group. This package accounts for genotyping errors and missing data and can estimate relatedness using any of seven different estimators (four non-likelihood-based and three likelihood-based). Using the *compareestimators* function, we generated simulated data from observed allele frequencies and assessed the performance of different non-likelihood estimators on the simulated data. We generated 200 simulated pairs of individuals for each degree of relatedness (i.e., half-sib, full-sib, parent–offspring, unrelated), and determined that the Wang (2002) estimator provided simulated estimates that best matched the observed data (see also Thrasher et al. 2018). Using the Wang (2002) estimator, we again generated 200 pairs of individuals for each degree of relatedness using the *familyism* function. The distributions generated from this function provided the bounds for assigning relationships when the values deviated from each predicted degree of relatedness. We then calculated pairwise relatedness between all individuals with the Wang (2002) estimator using the *coancestry* function (Wang 2011). These estimates were subsequently used to determine the genetic relationships between breeders and auxiliaries and between co-breeding individuals.

### Ethical note

All field methods were approved by the Cornell Institutional Animal Care and Use Committee (IACUC 2009-0105), Tulane University IACUC (2019-1715), and the James Cook University Animal Ethics Committee (A2100). The present study was permitted under Queensland Government Department of Environment and Heritage Protection Scientific Purposes Permit (WISP15212314). Our banding and blood sampling methods have been used previously in a closely related species with no observable negative effect (Webster *et al*. 2008).

## Results

### Group Size and Composition

From 2014 – 2017, we captured and sampled 858 individuals (319 adults and 539 nestlings). The number of social groups monitored ranged from 54 to 57 groups across all four years of the study (Table 1). Territories remained largely stable between years, but social groups changed due to demographic processes, so independent social groups were identified each year. In total, we identified and monitored 222 unique social groups during the study (Table 1, Figure 1).

**Table 1.** Number of Variegated Fairywren social groups, average group size, and composition by year.

| Year | No. of groups | Group size (mean ± SD) | No. of males (mean ± SD) | No. of females (mean ± SD) |
| --- | --- | --- | --- | --- |
| 2014 | 55 | 4.58 ± 1.55 | 2.84 ± 1.34 | 1.75 ± 0.62 |
| 2015 | 54 | 4.52 ± 1.50 | 2.70 ± 1.24 | 1.81 ± 0.52 |
| 2016 | 57 | 4.23 ± 1.76 | 2.44 ± 1.36 | 1.79 ± 0.73 |
| 2017 | 56 | 4.38 ± 1.74 | 2.50 ± 1.36 | 1.88 ± 0.81 |
| Mean | 55.5 | 4.42 ± 1.64 | 2.62 ± 1.33 | 1.81 ± 0.67 |

**Figure 1.**
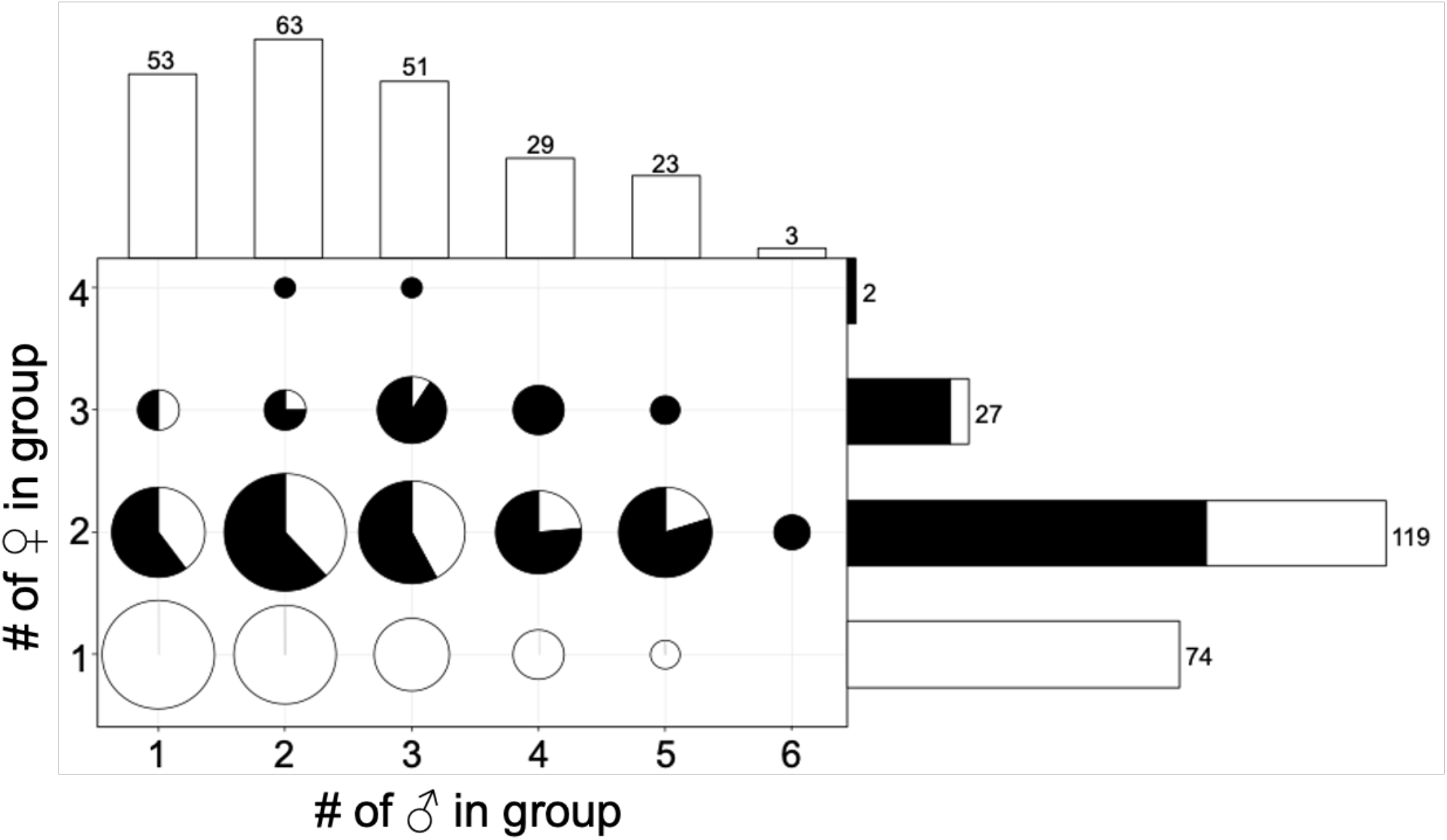
Composition of Variegated Fairywren social groups from 2014 – 2017 (N = 222 group years). The size of each circle indicates how common that composition was relative to the total number of groups. Histograms along both axes show the number of groups with a particular number of individuals of each sex. The black portion within circles and bars indicates the proportion of groups that had multiple breeding females, and white indicates the proportion of groups with a single breeding female.

In the 2016 field season, nearly all (95%) of dyadic relationships that occurred within routine monitoring groups were also present in the network-defined groups, (Figure 2). The few mismatches between these methods can be explained by a small number of groups: in one instance, the network analysis split a pair that was together in the field-defined groups; in another instance, a single bird was placed in a different group in the network analysis; and there were two instances where the network and database differed on whether to split a large group or to keep it together. Permutation analyses revealed that the network-defined groups were more structured than expected by chance (p<0.01, Figure S2).

**Figure 2.**
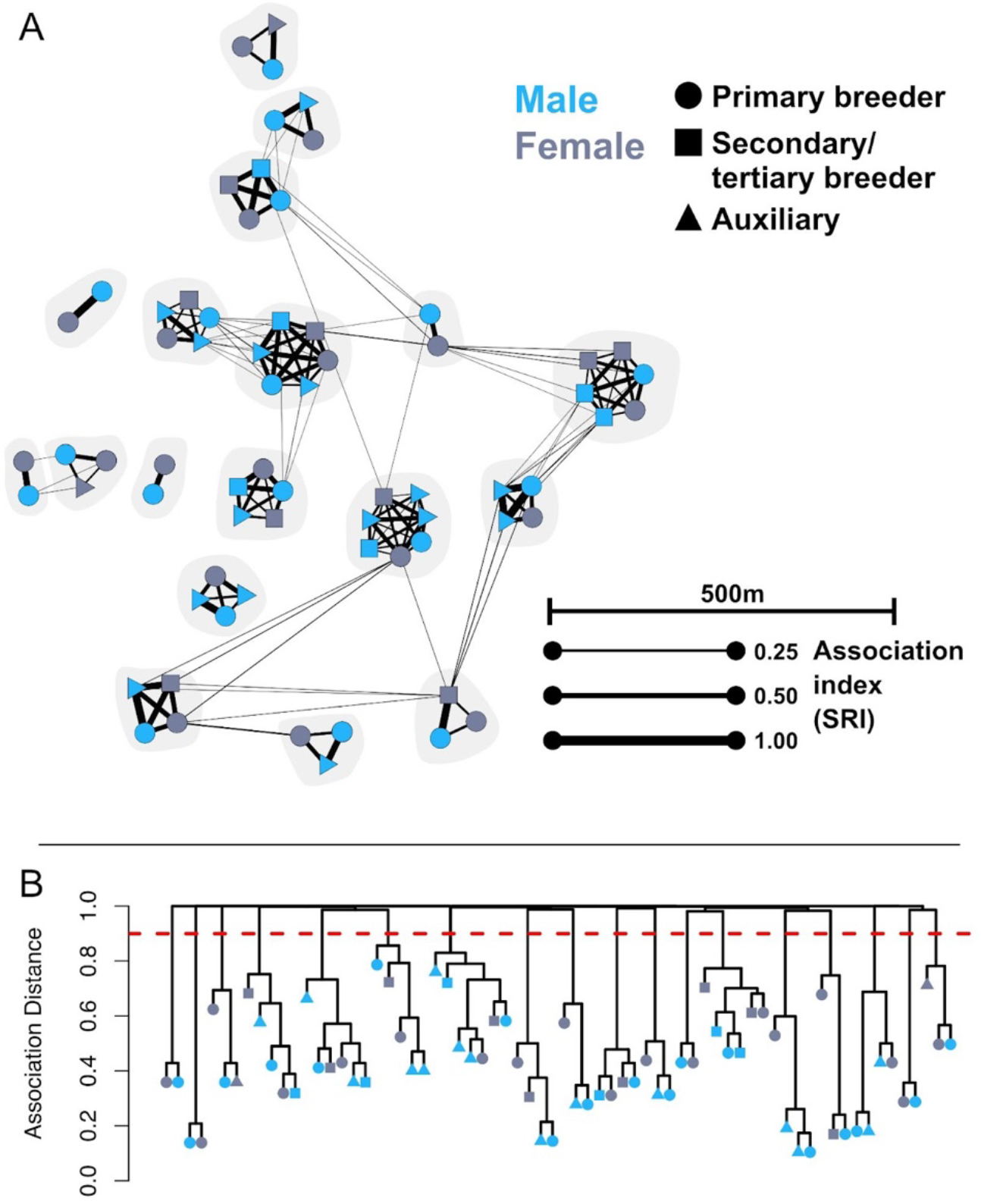
Variegated Fairywren social relationships and group structure during the breeding season. **A)** A representative subset of the social network from the 2016 breeding season. Each node represents an individual bird and lines connecting nodes are sized relative to how often those two individuals were seen together using the Simple Ratio Index (SRI). Thicker lines indicate individuals seen associating more often than thinner lines. Sex is represented by color, breeding status is represented by node shape, and social group membership is represented by shading behind nodes. Social groups are plotted geographically to where each group spent the most of its time. **B)** Dendrogram used to identify social group membership for the subset of individuals in the network above. Each node represents an individual bird as above and individuals connected by lines that do not cross the horizontal red line are considered in the same social group. Association distance (y-axis) is the inverse of the association index (1-SRI).

Social groups ranged in size from 2 – 8 adults (Table 1) and the adult sex ratio was male-biased (1.44:1). The number of males in a social group was more variable than was the number of females (Table 1, Figure 1), ranging from 1 – 6 males and 1 – 4 females per group. Of all social groups, 13% were socially monogamous pairs (n = 29), 40% were cooperative groups with one breeding female (n = 87), and 47% were cooperative groups with multiple co-breeding females (n = 105). Groups with multiple females were common (67%; n = 148; Figure 1), and over two-thirds of those exhibited plural breeding (i.e., multiple co-breeding females within a single social group). Of these plural breeding groups (n = 105), 90% had two breeding females and 10% had three breeding females.

### Origins and Statuses of Known Individuals

We followed 115 nestlings (74 males; 41 females) to adulthood and identified 60 yearling immigrants (19 males; 41 females) during our study (Table S1). In their first year, males were typically philopatric (89% of males) whereas most females dispersed from their natal territory as yearlings (59%; Table S1). Females that did not disperse as yearlings did so the following year, except one female that remained on her natal territory and became the primary breeding female in her fourth year after the disappearance of her mother. This was the only case of a female inheriting a breeding position on her natal territory during our study.

Sexes differed in their likelihood of becoming breeding adults. Of those hatched on territories within our study site (“local” females and males), most local females became breeders during either their first (37%) or second (24%) breeding seasons, and the remaining 39% spent their first season as a non-breeder and then disappeared (presumably dispersed or died); none of these females remained as a non-breeder for more than one season. In contrast, a large proportion (39%) of the local males hatched on the study site remained as non-breeding auxiliaries throughout the study, 20% became breeders in their first and 24% became breeders in their second breeding season. Only 16% of auxiliary males disappeared after spending their first season as a non-breeder. Immigrants were more likely to breed in their first breeding season on our study site in both males (52.6%) and females (65.9%).

### Relatedness within Social Groups

Average relatedness of adults within social groups was 0.11 (SD ± 0.21), suggesting a mix of closely related and unrelated individuals. Within a social group, pairwise relatedness estimates were generally much higher between males than between females (males: mean ± SD = 0.18 ± 0.20, n = 495; females: 0.06 ± 0.20, n = 147), suggesting that many females were likely immigrants to the group and unrelated to one another.

We further investigated relationships among group members by assessing patterns of pairwise relatedness between different categories of group members (Figure 4). Co-breeding females were mostly unrelated to each other (mean ± SD = −0.01 ± 0.07; Figure 3a), as would be expected if breeding females were mostly immigrants from other social groups, which is supported by our behavioral observations (see above). Pairwise comparisons between breeding and non-breeding females revealed a bimodal distribution (Figure 3b), one mode of highly related individuals (*r*-value ~ 0.50), as expected of mother-daughter or sister pairs, and the other of unrelated individuals (*r*-value ~ 0.0). The distribution of relatedness of males within social groups was more continuous due to individuals of varying degrees of relatedness between the expected values for unrelated individuals and full sibs (r = 0.0 and 0.5, respectively; Figure 3d).

**Figure 3.**
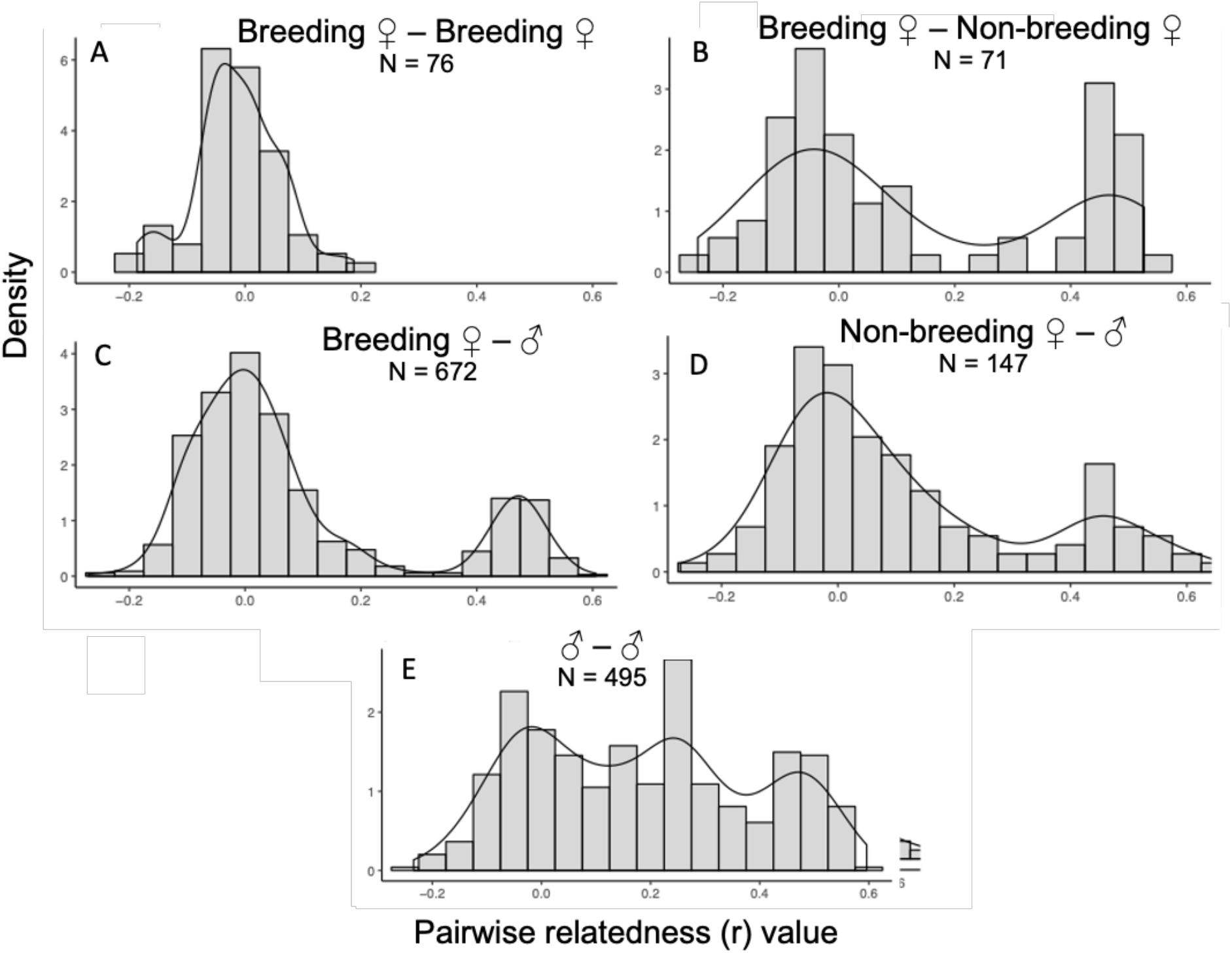
Distributions of pairwise genetic relationships of Variegated Fairywren social group members based on sex and breeding status of females. Comparisons include those between A) breeding females, B) breeding females and non-breeding females, C) breeding females and all males, D) non-breeding females and non-breeding males, and E) all males. Note varying y-axes across panels.

Pairwise relatedness between breeding females and all males within a social group (Figure 3c) was similar to that of breeding females and non-breeding females (Figure 3b), but with fewer relatives. The distribution of pairwise relatedness between non-breeding females and all males also exhibited a bimodal distribution of unrelated individuals and highly related individuals, but the majority were predominantly unrelated (Figure 3e).

## Discussion

Our study reaffirms the Variegated Fairywren as a cooperative breeder (Buchanan and Cockburn 2013; Mclean *et al*. 2017b; Mclean *et al*. 2017a), but also shows that the breeding system involves highly complex patterns of relatedness and reproduction within social groups. Although some groups were socially monogamous pairs that reared and fledged offspring in the absence of auxiliary helpers, most groups (nearly 87%) were cooperative groups with auxiliary adults. Cooperative groups were extremely variable in size and composition, ranging widely in number of males and females occupying breeding or auxiliary roles (Fig. 1). Although most groups were male-biased, well over half of all social groups contained multiple females, and of those a large majority (71%) were groups that contained two or more breeding females. Multi-female groups in our population formed as the result of recruitment of daughters on their natal territory, the arrival of immigrant females to a group with an established breeding female, or combinations of the two. Plural breeding groups always manifested through immigrant females joining groups with established breeding females. This pattern was confirmed by our genetic analysis, which showed that co-breeding females were never close relatives.

### Social Network Analysis of Group Membership

Fairywrens exhibit unusually high levels of extra-pair paternity (EPP) with both males and females embarking on off-territory forays in search of extrapair matings (Rowley and Russell 1990; Double and Cockburn 2000; Potticary et al. 2016; Leitão et al. 2019; Boersma et al. 2022). Connections between social groups in our study were likely the product of individuals engaging in extra-pair courtship (Fig. 2a). A few groups were isolated from the rest of the population, likely due to geographic boundaries (Welklin *et al*. 2023). Ultimately, the groups identified via social network analysis in 2016 matched the groupings assigned through routine monitoring almost exactly (95%), thus lending confidence in our group assignments across study years.

### Relatedness within Social Groups

One of the most striking patterns revealed by our genetic analysis was that plurally breeding females within a group were always unrelated to one another. While plural breeding has been noted in other Fairywrens, co-breeding females in those species were typically close relatives (i.e. mothers and daughters; Rowley et al. 1989; Russell and Rowley 2000). In other species, such as the Galapagos Mockingbird (*Mimus parvulus*; Curry 1988) and Mexican Jay (*Aphelocoma wollweberi*; Barkan *et al*. 1986; Li and Brown 2000), plural breeding by close relatives is common and is usually the result of limited breeding opportunities outside of the social group. Secondary breeding females in these species often initiate their own nests, but generally at a lower level than primary breeders. In our study plural breeders were always non-relatives, which is uncommon in birds (Riehl 2013), and the first published evidence in Fairywrens. Co-breeding females jointly defend territories with other members of the social group, so could benefit from enhanced capacity to defend limited resources and deter predators and brood parasites (Riehl and Jara 2009; Feeney *et al*. 2013; Shen *et al*. 2017b). Determining the fitness consequences for plural breeding will be informative to our understanding of mating systems and the evolution of cooperation.

Relatedness patterns of males within social groups was a mixture of nonrelatives (r=0), moderate relatives (r=0.25), and close relatives (r=0.5). Varying levels of relatedness among males in the same social group is likely the product of high rates of EPP commonly observed in Fairywrens (Dunn *et al*. 1995; Webster *et al*. 2004; Johnson and Pruett-Jones 2018), and in rarer cases, immigrant males joining groups. The extent to which offspring are related to breeding pairs can vary across cooperative breeders, as does their help with provisioning and whether they attempt their own reproduction within the group (Williams 2004; Riehl and Jara 2009; Raihani *et al*. 2010; Groenewoud *et al*. 2018). Consistent with other Fairywren species, greater female dispersal led to a strong male sex bias among auxiliaries in our study (Russell and Rowley 2000; Webster *et al*. 2004; Potticary *et al*. 2016; Johnson and Pruett-Jones 2018; Leitão *et al*. 2019).

Given the complexity of relatedness within social groups, it is likely that cooperation among adults is maintained via both indirect and direct fitness benefits (West *et al*. 2007). For both sexes, some individuals remain on their natal territories as non-breeding helpers, likely deriving some kin-selected benefits (West *et al*. 2007; Kingma *et al*. 2010; Bourke 2014), though direct benefits are also possible. This strategy is common in males, but much less so for females, with few females remaining as non-breeding helpers beyond their first year. In addition, immigrants of both sexes joined established social groups as unrelated auxiliaries. Most immigrants eventually adopted a breeding role in the group they joined, suggesting that non-kin auxiliaries are immigrants that join the group to queue for breeding opportunities. Unrelated male auxiliaries may also benefit by sneaking copulations with females (Riehl 2013). Future work in this system can resolve the selective pressures underlying complex sociality.

### Conclusions

Our findings indicate that Variegated Fairywrens exhibit a complex social system characterized by a dynamic combination of breeding and non-breeding individuals of varying relatedness. This species is also unique among *Malurus* Fairywrens in that unrelated co-breeding females are often present in the same social group. The complexity of this social system offers an ideal opportunity to answer questions about cooperation and conflict in social groups. Although tradeoffs are likely present for all group members, it is particularly important for follow-up work to focus on unrelated plurally-breeding females given the likely absence of any kin-selected benefits. Co-breeding females do cooperate in territory defense, and thus might derive benefits from enhanced protection from conspecifics, predators, and brood parasites. Plural breeding may also incentivize cooperation in males as opportunities for breeding are more likely as the number of breeding females increases. Continued work in this system could provide important insights into the evolution of cooperative behavior among diverse taxa, particularly which factors incentivize cooperation among individuals of varying relatedness.

## Acknowledgements

We sincerely thank the many field technicians that contributed to our intensive field monitoring efforts: R. Bracken, C. Brock, J.D. Brooks, L. Dargis, Z. Davis, B. Donnelly, C. Egan, M. Freeby, L. Fried, K. Gielow, M.M. Gray, J. Hiciano, H. Kraus, A. Miller, R. Neil, J. Platzer, P. Queller, H. Short, A. Werrell, and E. Zarri, Members of the Webster lab and William Feeney provided valuable input. We also thank the Lovette Lab at the Cornell Lab of Ornithology, and especially B. Butcher, for assistance and guidance with molecular methods. Thanks to the National Science Foundation (USA), the Cornell Lab of Ornithology, and the department of Neurobiology and Behavior at Cornell University for supporting this work.

## Supplementary Materials

**Table S1.** Origins of known-age yearling adult Variegated Fairywrens. Immigrants are divided into two categories within sex: individuals new to the population (UNK) and individuals born in the population that dispersed from one social group to another (KN).

|  | 2014 | 2015 | 2016 | 2017 | Total |
| --- | --- | --- | --- | --- | --- |
| **Natal Female** | 2 | 8 | 0 | 7 | 17 |
| **Natal Male** | 14 | 17 | 13 | 21 | 65 |
| **Imm. Female - KN** | 6 | 6 | 6 | 6 | 24 |
| **Imm. Male - KN** | 0 | 6 | 0 | 3 | 9 |
| **Imm. Female - UNK** | - | 7 | 17 | 17 | 41 |
| **Imm. Male - UNK** | - | 6 | 9 | 4 | 19 |
| **Total** | **22** | **50** | **45** | **58** | **175** |

**Table S2.**
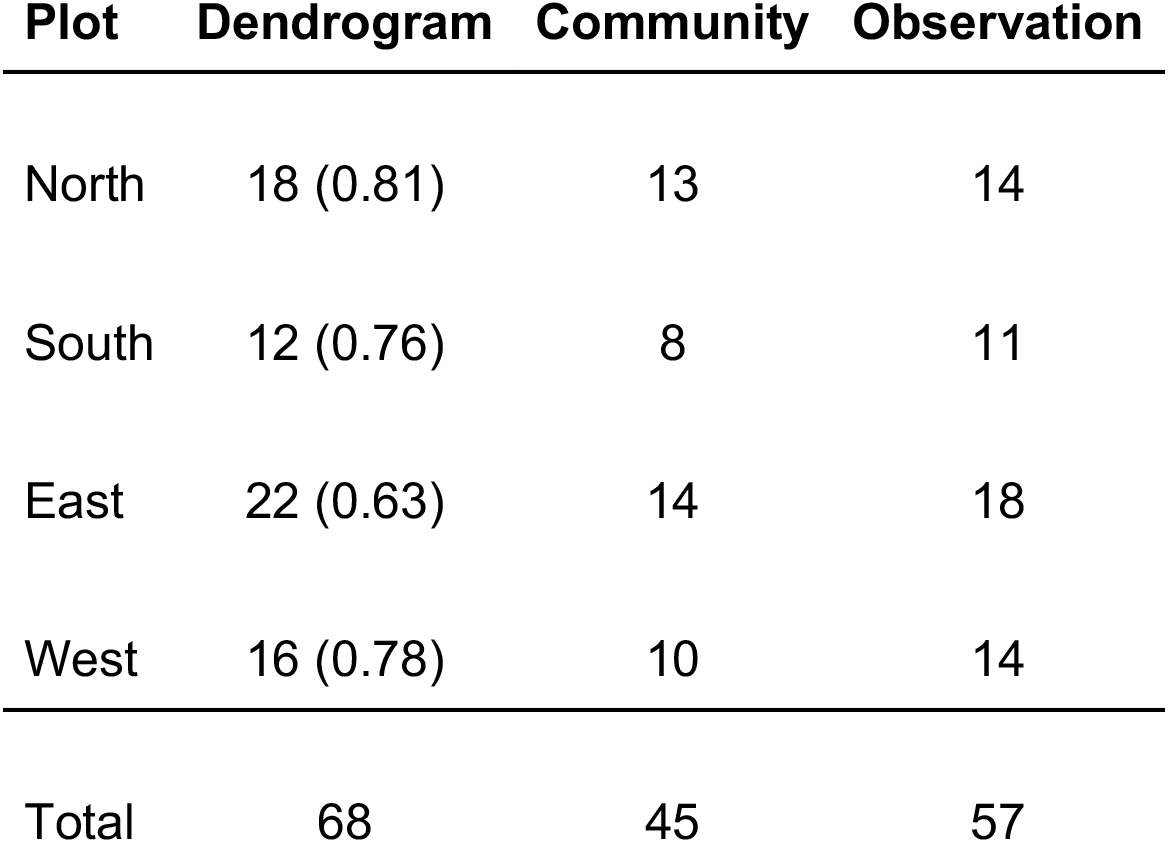
Comparison of calculated number of Variegated Fairywren groups in population for 2016 breeding season calculated by different methods: silhouette/ dendrogram, network communities, and by field observation assignment. Cutoff silhouette values are indicated in parentheses for the dendrogram method.

**Figure S2.**
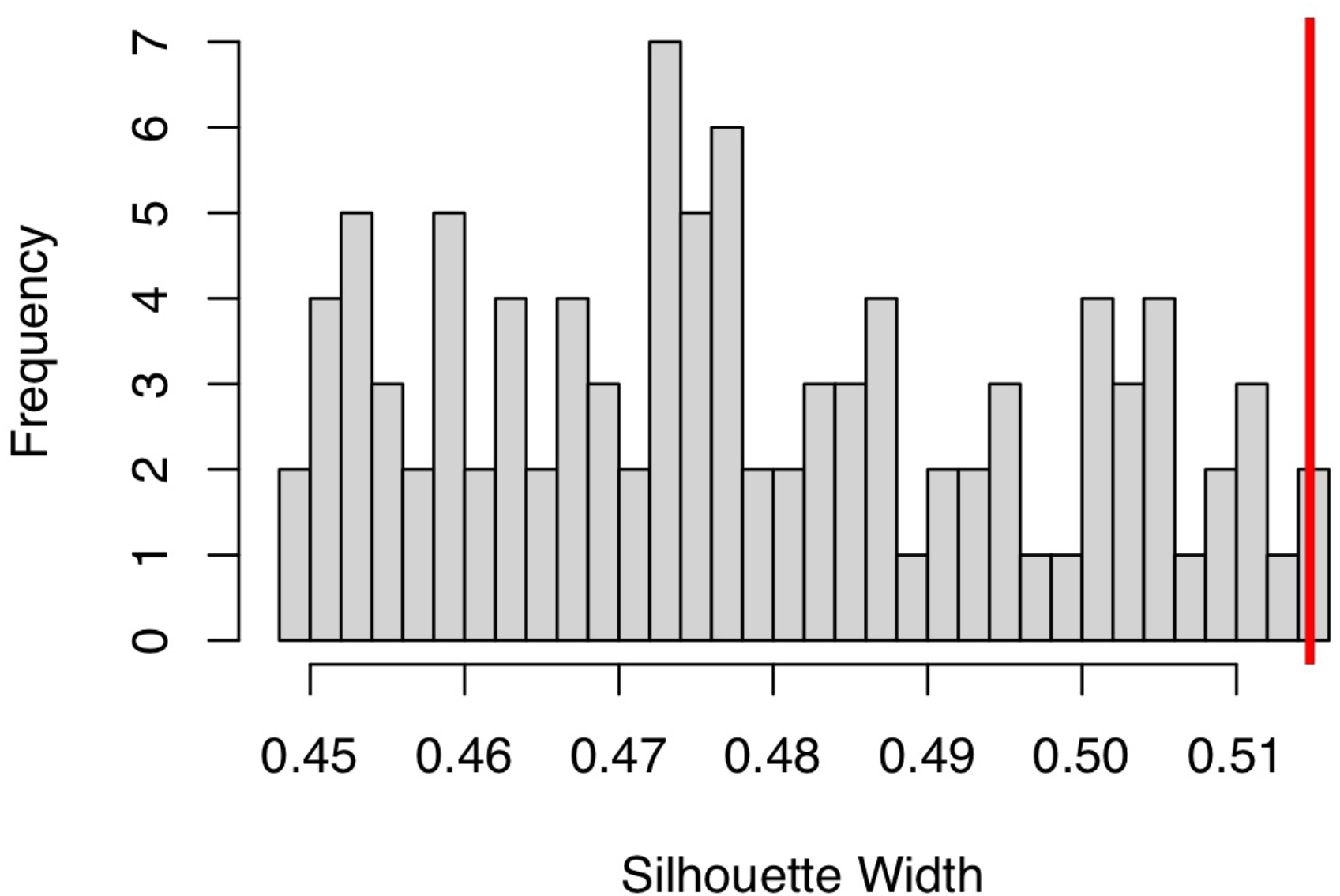
Histogram showing the observed silhouette width of the 2016 social network (vertical red line) to those of 100 randomized social networks (gray bars). The observed silhouette width was greater than the silhouette widths of each randomized network indicating the observed network was more structured than expected by chance.

## Notes

### Competing Interest Statement

The authors have declared no competing interest.

